# *USH2A* High-Capacity Adenoviral Vector Restores usherin, whirlin, and ADGRV1 in Photoreceptor Cells of Retinal Organoids

**DOI:** 10.64898/2026.09.14.751468

**Authors:** Rossella Valenzano, Xuefei Lu, Aat A. Mulder, Elon H.C. van Dijk, Roman I. Koning, Manuel A.F.V. Gonçalves, Jan Wijnholds

## Abstract

Development of gene therapies for *USH2A*-associated diseases has been limited by the large size of the *USH2A* coding sequence, which exceeds the packaging capacity of adeno-associated viral vectors. Here, we evaluated high-capacity adenoviral vectors (HC-AdVs) as vehicles for a mutation-independent *USH2A* gene supplementation strategy in *USH2A* human retinal organoids. First, we generated human induced pluripotent stem cell (hiPSC) lines carrying a nonsense mutation in *USH2A* exon-61 (61KO). Similarly to our previously established exon-13 knockout model (13KO), 61KO hiPSC-derived retinal organoids showed reduced levels of usherin and its interacting proteins, adhesion G protein-coupled receptor V1 (ADGRV1), and whirlin, at the photoreceptor connecting cilium. Delivery of a HC-AdV5 encoding full-length *USH2A* under the control of the photoreceptor-specific human rhodopsin kinase (hGRK1) promoter restored usherin expression and promoted re-localization of ADGRV1 and whirlin to their physiological sites at the connecting cilium in 13KO and 61KO photoreceptors. Unexpectedly, usherin expression was also observed within mutant Müller glial cells. These findings provide proof-of-concept that HC-AdV5-mediated delivery of full-length *USH2A* can successfully promote the rescue of usherin and relocate its interacting partners in *USH2A*-defective retinal organoids, demonstrating its potential as a variant-independent therapeutic approach for inherited retinal disorders caused by *USH2A* mutations.

## Introduction

Pathogenic variants of *USH2A* are among the leading cause of inherited retinal degeneration, yet the large size of its coding DNA sequence (CDS) has limited the development of broadly applicable therapies.^1, 2^ Here, we investigated whether high-capacity adenoviral vector-mediated delivery of the full-length *USH2A* CDS under the control of a photoreceptor-specific gene promoter restores usherin expression and rescues the localization of ADGRV1 and whirlin at the photoreceptor connecting cilium (CC) in two models of *USH2A* mutant retinal organoids.

Usher Syndrome type II is an autosomal recessive disorder that manifests vision loss accompanied by hearing dysfunction. While *USH2A* exon-13 harbors three of the top five most recurrent variants, namely c.2299delG, c.2276G>T, and c.2802T>G,^2^ several additional mutational hotspots have been identified, including in exon-61, where the c.11864G>A variant is predominantly found in homozygous state.^3, 4, 5^ A knock-in mouse carrying the human c.2299delG variant developed retinal degeneration, characterized by increased cellular stress and mislocalization of usherin and its interacting partners, ADGRV1 and whirlin, and exhibited hearing loss due to disrupted stereocilia bundles.^6, 7^ In *Ush2a*-ΔEx12 mice, skipping of the human *USH2A* exon-13 generated a shortened usherin that restored ciliogenesis and hearing, corrected cone opsin localization, and reduced gliosis.^8^ Human induced pluripotent stem cell (hiPSC)-derived retinal organoids generated from patient lines carrying compound c.2299delG and c.1256G>T *USH2A* variants exhibited thinning of the outer nuclear layer, with rod-specific structural loss, and progressive degeneration of photoreceptor inner and outer segments.^9^ Moreover, distinct *USH2A* variants associated with either retinitis pigmentosa or Usher Syndrome produced different retinal phenotypes in organoids.^10^ In our previous study, introduction of a premature stop codon in exon-13 of *USH2A* resulted in loss of usherin long isoform B, ADGRV1, and whirlin from the photoreceptor connecting cilium, accompanied by transcriptional changes in Müller glial cells (MGCs) suggestive of homeostasis disruption and activation of the innate immune response.^11^

The large 15.6-kb *USH2A* CDS represents a major obstacle for the development of adeno-associated viral (AAV) vector-based gene therapy, since it exceeds the ∼4.7-kb packaging capacity of AAV capsids. Consequently, current therapeutic approaches have focused primarily on CRISPR-dependent correction of recurrent pathogenic variants.^12^ For instance, base editing of the c.11864G>A variant restored usherin expression in more than 50% of transduced cells in preclinical studies.^13^ While promising, these strategies remain variant-specific and therefore cannot address the extensive allelic heterogeneity of *USH2A*.

High-capacity adenoviral (HC-AdV) vectors offer a variant-independent alternative. With a packaging cargo of up to 36-kb, HC-AdVs can accommodate the full-length *USH2A* CDS and associated regulatory elements to potentially mediate long-term transgene expression in post-mitotic cells from persisting vector episomes.^14, 15^ HC-AdVs have demonstrated efficient gene delivery in liver, neurons, endothelium, and skeletal muscle, where, in the latter tissue, they enabled restoration of dystrophin upon the delivery of the full-length 11.1-kb *DMD* CDS together with regulatory sequences.^16–19^ In the retina, HC-AdVs have mediated sustained *EGFP* expression in the rat retinal pigment epithelium, without significant toxicity or inflammatory responses.^15^ Finally, in both mice and human iPSC-derived retinal organoids, HC-AdV5 and fiber-modified HC-AdV5.F50 demonstrated efficient transduction of photoreceptors and Müller glial cells, supporting their potential as vectors for the delivery of large retinal disease-causing genes.^20, 21, 22^

## Results

### Generation and characterization of CRISPR/Cas9-engineered USH2A 61KO hiPSC-derived retinal organoids

To obtain a second independent model of *USH2A* loss-of-function for testing *USH2A* HC-AdV5 gene supplementation besides our previously characterized *USH2A* 13KO organoids,^11^ here we targeted another mutational hotspot within the 3ʹ region of *USH2A*. Using a CRISPR/Cas9-based approach, we introduced a CATG sequence in *USH2A* exon-61 (c.11992_11993insCATG) of the parental LUMC0004iCTRL10 hiPSC line,^23^ here renamed as ISO-CTRL. This modification causes a frameshift with the formation of a premature stop codon, resulting in a predicted truncated protein, p.(Tyr3998Serfs*2), and creates a recognition site for the restriction enzyme BspHI for screening of edited clones (Figure S1A). The mutant p.(Tyr3998Serfs*2) resembles in length and domain structure shorter usherin variants found in patients and annotated in the LOVD database (https://www.lovd.nl/), particularly those affecting exons 60-62 (Figure 1), where deletions or missense mutations similarly lead to truncation of usherin within the C-terminal fibronectin type III domains.

**Figure 1.**
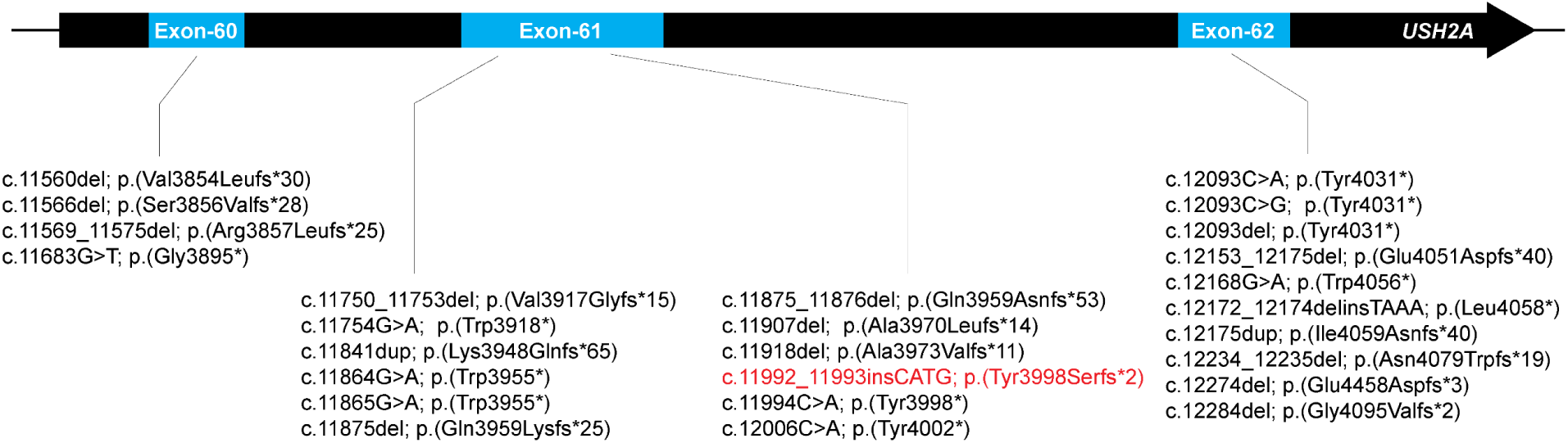
List of *USH2A* pathogenic variants locating in exons 60-62 and resulting in a predicted truncated usherin. The *USH2A* c.11992_11993insCATG; p.(Tyr3998Serfs*2) presented in this study (annotated in red) recapitulates the structure of usherin as predicted from other pathogenic variants in exons 60, 61, and 62, leading to premature stop codons.

Following electroporation of the ISO-CTRL line with the targeted gene editing machinery consisting of *USH2A*-specific Cas9:gRNA complexes and a single-stranded oligonucleotide donor template (Figure S1A), hiPSC subclones were screened by PCR and restriction fragment length analysis. Clones showing digestion with the BspHI enzyme were labeled as *USH2A* 61KO H6, 61KO G11, and 61KO E9, and the correct CATG insertion was confirmed by Sanger sequencing (Figure S1B). Importantly, recurrent copy number variations were not detected (Figure S1C) and G-banding karyotyping revealed normal metaphases (Figure S1D). All 61KO subclones exhibited short tandem repeat profiles consistent with the ISO-CTRL line (Figure S1E). To further complete the molecular characterization of these new *USH2A* mutant hiPSC lines, off-target mutagenesis analysis was performed on the top five predicted loci identified using the online tools provided by Integrated DNA Technologies (IDT) (https://eu.idtdna.com/). Sanger sequencing chromatograms showed no abnormal peaks in the regions surrounding the candidate off-target sites (Figure S2).

*USH2A* 61KO H6, G11, and E9 hiPSC subclones were then differentiated into *USH2A* mutant retinal organoids along with their ISO-CTRL parental line. We observed comparable morphology between ISO-CTRL and *USH2A* 61KO organoids at differentiation day (DD) 225, with the typical brush border marking the photoreceptor layer (Figure 2A). The distribution of the rod marker rhodopsin between the photoreceptor segments and the outer nuclear layer (ONL) followed comparable patterns in both control and mutant organoids (Figure 2B, 2C). ARL13B-marked ciliary length and the outer nuclear layer thickness showed no statistically significant differences in any of the 61KO subclones when compared to ISO-CTRL (Figure 2D, 2E).

**Figure 2.**
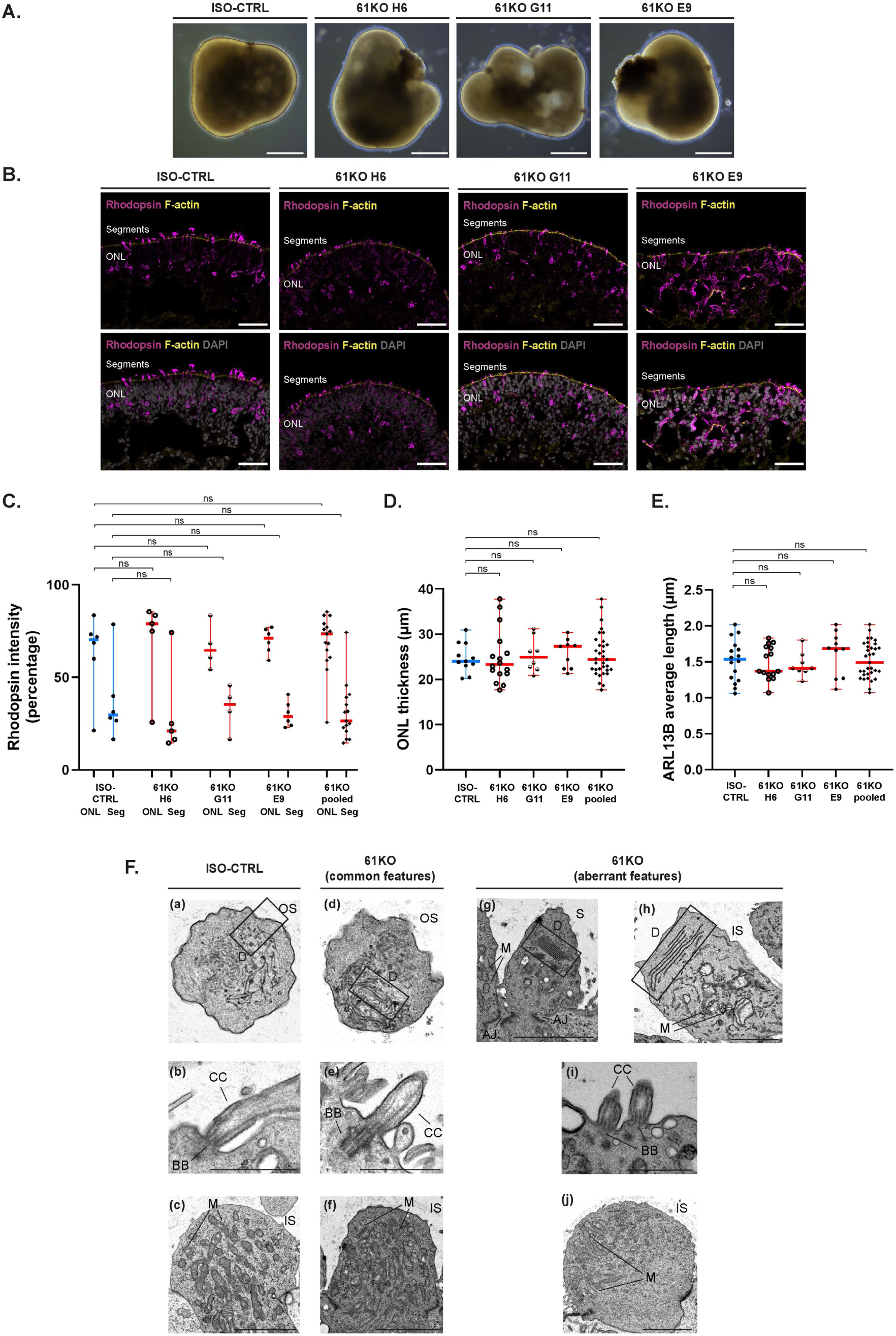
Phenotypic analysis of the photoreceptor layer of *USH2A* 61KO retinal organoids at DD225. (A) Brightfield images of ISO-CTRL, *USH2A* 61KO H6, 61KO G11, and 61KO E9 retinal organoids at DD225. Scale bars: 200 µm. (B) Representative immunofluorescence images of rhodopsin (magenta) and F-actin (yellow) in ISO-CTRL and 61KO retinal organoids with and without DAPI (grey). Scale bars: 50 µm. (C) Quantitative analysis of rhodopsin intensity distribution between the outer nuclear layer (ONL) and the photoreceptor segments (Seg) with and without DAPI (grey). Data are presented as median with range. ISO-CTRL ONL: median, 29.7; minimum, 16.5; maximum, 78.69; ISO-CTRL Seg: median, 70.31; minimum, 21.31; maximum, 83.5; 61KO H6 ONL: median, 20.92; minimum, 14.56; maximum, 74.31; 61KO H6 Seg: median, 79.08; minimum, 25.69; maximum, 85.44; 61KO G11 ONL: median, 35.39; minimum, 16.53; maximum, 45.68; 61KO G11 Seg: median, 64.61; minimum, 54.32; maximum, 83.47; 61KO E9 ONL: median, 28.82; minimum, 22.84; maximum, 40.83; 61KO E9 Seg: median, 71.19; minimum, 59.17; maximum, 77.16; 61KO pooled ONL: median, 26.46; minimum, 14.56; maximum, 74.31; 61KO pooled Seg: median, 73.54; minimum, 25.69; maximum, 85.44. Number of organoids used: ISO-CTRL, n = 6; 61KO H6, n = 6, 61KO G11, n = 4, 61KO E9, n = 6, 61KO pooled, n = 15, from three differentiations. Significance was calculated by Kruskal-Wallis followed by Dunn’s multiple comparisons test and indicated as ns (not significant). Statistical analysis: *p* = 0.6317, *p* > 0.9999, *p* > 0.9999 and *p* > 0.9999, from left to right. (D) Quantitative analysis of ONL thickness. Data are presented as median with range. ISO-CTRL: median, 24.01; minimum, 20.20; maximum, 30.96; 61KO H6: median, 23.31; minimum, 17.69; maximum, 37.77; 61KO G11: median, 24.92; minimum, 20.89; maximum, 31.22; 61KO E9: median, 27.33; minimum, 21.29; maximum, 30.39; 61KO pooled: median, 24.40; minimum, 17.69; maximum, 37.77. Number of organoids used: ISO-CTRL, n = 11; 61KO H6, n = 16, 61KO G11, n = 8, 61KO E9, n = 9, 61KO pooled, n = 33, from three differentiations. Significance was calculated by Kruskal-Wallis followed by Dunn’s multiple comparisons test and indicated as ns (not significant). Statistical analysis: *p* > 0.9999 for all comparisons from left to right. (E) Quantitative analysis of ARL13B average length. Data are presented as median with range. ISO-CTRL: median, 1.535; minimum, 1.060; maximum, 2.020; 61KO H6: median, 1.370; minimum, 1.070; maximum, 1.830; 61KO G11: median, 1.410; minimum, 1.230; maximum, 1.800; 61KO E9: median, 1.685; minimum, 1.070; maximum, 2.020; 61KO pooled: median, 1.490; minimum, 1.070; maximum, 2.020. Number of organoids used: ISO-CTRL, n = 16; 61KO H6, n = 15, 61KO G11, n = 8, 61KO E9, n = 10, 61KO pooled, n = 33, from three differentiations. Significance was calculated by Kruskal-Wallis followed by Dunn’s multiple comparisons test and indicated as ns (not significant). Statistical analysis: *p* = 0.9920, *p* = 0.9654, *p* = 0.7056 and *p* > 0.9999, from left to right. (F) Representative transmission electron microscopy images of photoreceptors in ISO-CTRL and 61KO retinal organoids at DD225. Images show common features as outer segment (OS)-like structures with immature disc membranes (D), outlined by black boxes (a, d), the presence of a basal body (BB) and connecting cilium (CC) (b-e), and inner segment (IS)-like structures enriched with mitochondria (M) (c-f). Images in F(g-j) show aberrant features observed in the 61KO organoids, including stacked membranous discs (D) within a photoreceptor segment (S) connected via adherens junction (AJ) to a mitochondria (M)-enriched photoreceptor inner segment (g), stacked membranous discs (D) within a photoreceptor inner segment (IS) presenting few mitochondria (M) (h), double basal bodies (BB) and connecting cilia (CC) within a single IS (i), and polarized distribution of mitochondria (M) within the IS (j). Discs (D) in F(g-h) are outlined by black boxes. Scale bars in F(a, b, d, e, g, i) are 1 µm; scale bars in F(c, f, j) are 3 µm; scale bar in F(h) is 2 µm. Number of organoids imaged: ISO-CTRL *n* = 1, 61KO H6 *n* = 2, 61KO G11 *n* = 2, and 61KO E9 *n* = 2, from one differentiation.

Transmission electron microscopy analysis revealed that *USH2A* 61KO organoids at DD225 retained key features of photoreceptors comparable to those observed in the ISO-CTRL sample, including mitochondria enriched within inner-segment (IS)-like structures, a connecting cilium extending from a basal body, and immature discs within outer-segment (OS)-like structures (Figure 2F(a-f)). In addition to these preserved common features, occasional ultrastructural abnormalities were observed. These included stacked membranous discs within an aberrant photoreceptor segment adjacent to a mitochondria-enriched IS-like structure, discs within an abnormal IS-like region containing few mitochondria, the presence of two basal bodies and corresponding connecting cilia within a single inner segment, and an apical redistribution of mitochondria within IS-like regions (Figure 2F(g-j)). Owing to the limited number of organoids examined, it remains unclear whether these ultrastructural abnormalities represent a reproducible phenotype of the 61KO organoids or reflect variations within this differentiation batch.

### High-capacity adenoviral vectors as vehicles for USH2A gene supplementation therapy

Having established *USH2A* 61KO organoids as new disease models complementary to the previously characterized 13KO organoids, we next used them as human platforms to test *USH2A* gene supplementation strategies based on HC-AdV gene delivery.

We successfully generated a 33.3-kb plasmid carrying a 30.3-kb HC-AdV DNA molecular clone expressing the full-length human usherin protein. This plasmid was used to generate HC-AdV type-5 (HC-AdV5) vector particles, hereafter referred to as *USH2A* HC-AdV5 (Figure 3A). These viral particles have a total adenoviral vector cargo of 30.3-kb and express the 15.6-kb codon-optimized *USH2A* CDS under the control of the human rhodopsin kinase gene (hGRK1) promoter linked to the simian virus 40 small intron (SV40i) to drive selective expression of usherin in rod and cone photoreceptors. Moreover, a 3xHA tag was introduced at the C-terminal end of the usherin protein CDS to facilitate its detection and a woodchuck hepatitis virus post-transcriptional regulatory element (WPRE3) was placed downstream of the transgene, proximal to the herpes simplex virus (HSV) thymidine kinase polyadenylation signal, to increase expression at the post-transcriptional level. An additional HC-AdV5 vector carrying an *EGFP* transgene encoding the enhanced green fluorescent protein under the control of the ubiquitous CMV promoter, *EGFP* HC-AdV5, was used as control (Figure 3A). A similar *EGFP* HC-AdV5 previously showed efficient transduction of Müller glial cells and photoreceptors.^22^

**Figure 3.**
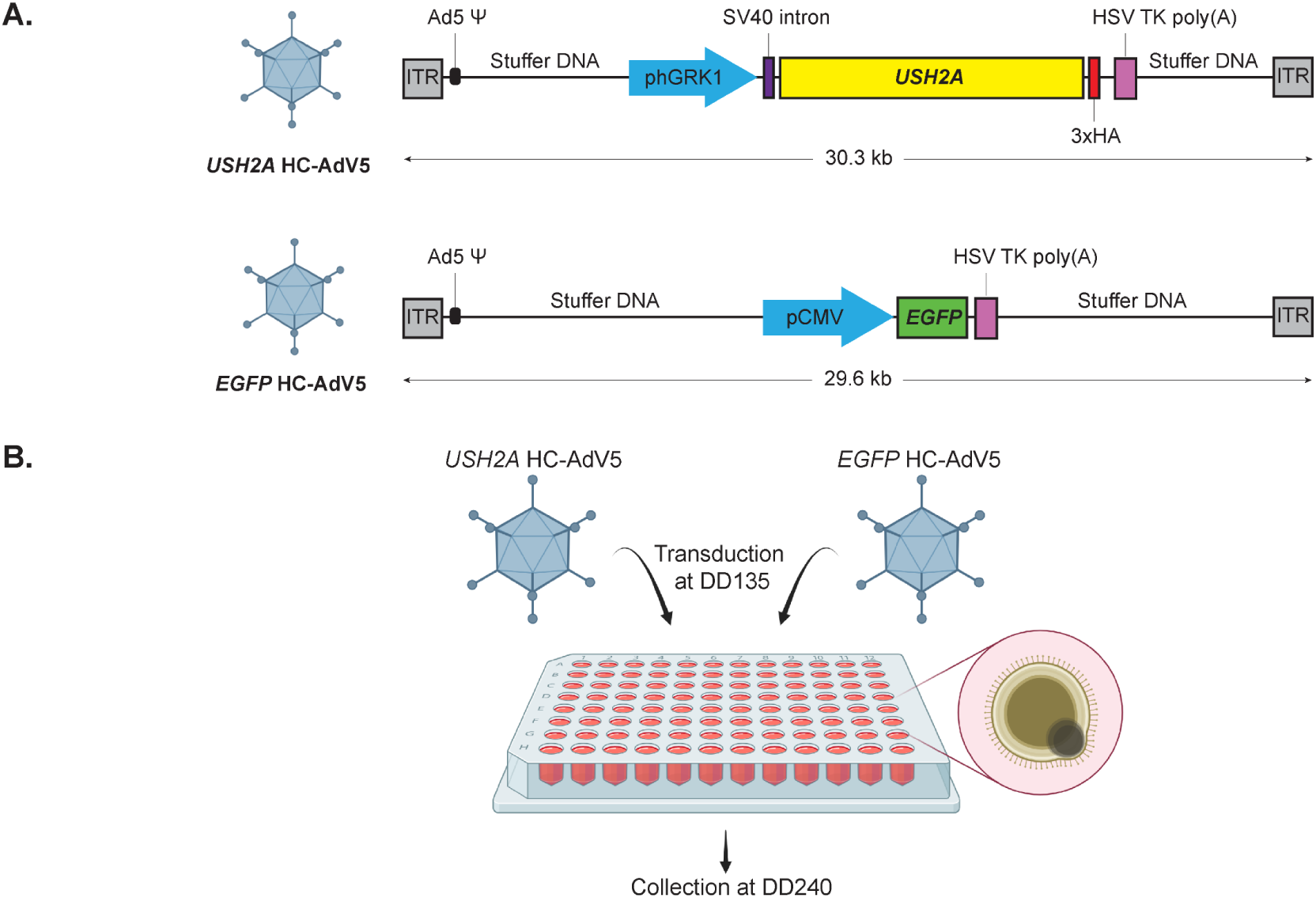
Transduction of retinal organoids with high-capacity adenoviral vectors. (A) Schematics of the *USH2A* HC-AdV5 and *EGFP* HC-AdV5 vectors. ITR and Ad5 Ψ, inverted terminal repeat and packaging signal for human adenovirus type 5, respectively. phGRK1, human G protein-coupled receptor kinase 1 promoter. HSV TK poly(A), herpes simplex virus thymidine kinase polyadenylation signal. (B) Representation of the timeline for *USH2A* HC-AdV5 transduction and collection of *USH2A* 13KO and 61KO organoids. Created in https://BioRender.com.

At DD135, *USH2A* 13KO and *USH2A* 61KO hiPSC-derived retinal organoids were transduced for 8 hours with *USH2A* HC-AdV5 together with *EGFP* HC-AdV5 (Figure 3B). To ensure a reliable assessment of the gene supplementation effect, only organoids with high morphological quality were included in the experiment. Therefore, 13KO subclones B7 and G8 and 61KO subclones H6 and G11 were selected for vector transduction, whereas 13KO E11 and 61KO E9 were excluded, since they did not generate retinal organoids of sufficient quality in this differentiation batch.

The *USH2A* 13KO B7, 13KO G8, and 61KO H6, and 61KO G11 organoids received a standardized total dose of 4.5 × 10^8^ vector particles (vp) per organoid, consisting of 4.1 × 10^8^ vp of *USH2A* HC-AdV5 and 0.4 × 10^8^ vp of *EGFP* HC-AdV5. HC-AdV5-treated and untreated *USH2A* 13KO and 61KO retinal organoids were subsequently collected at DD240 for immunofluorescence microscopy analysis, where ARL13B was employed as a ciliary marker to assess whether usherin expression was restored and correctly localized to the connecting cilium.

### USH2A HC-AdV5-transduced USH2A 13KO and 61KO retinal organoids show restoration of usherin, whirlin, and ADGRV1 at the photoreceptor connecting cilium

The analysis revealed that the usherin signal, which was below detection levels at the connecting cilium in both 13KO (B7 and G8) (Figure 4A) and 61KO (H6 and G11) (Figure 4B) untreated organoids, was detectable in proximity to ARL13B upon *USH2A* HC-AdV5 treatment.

**Figure 4.**
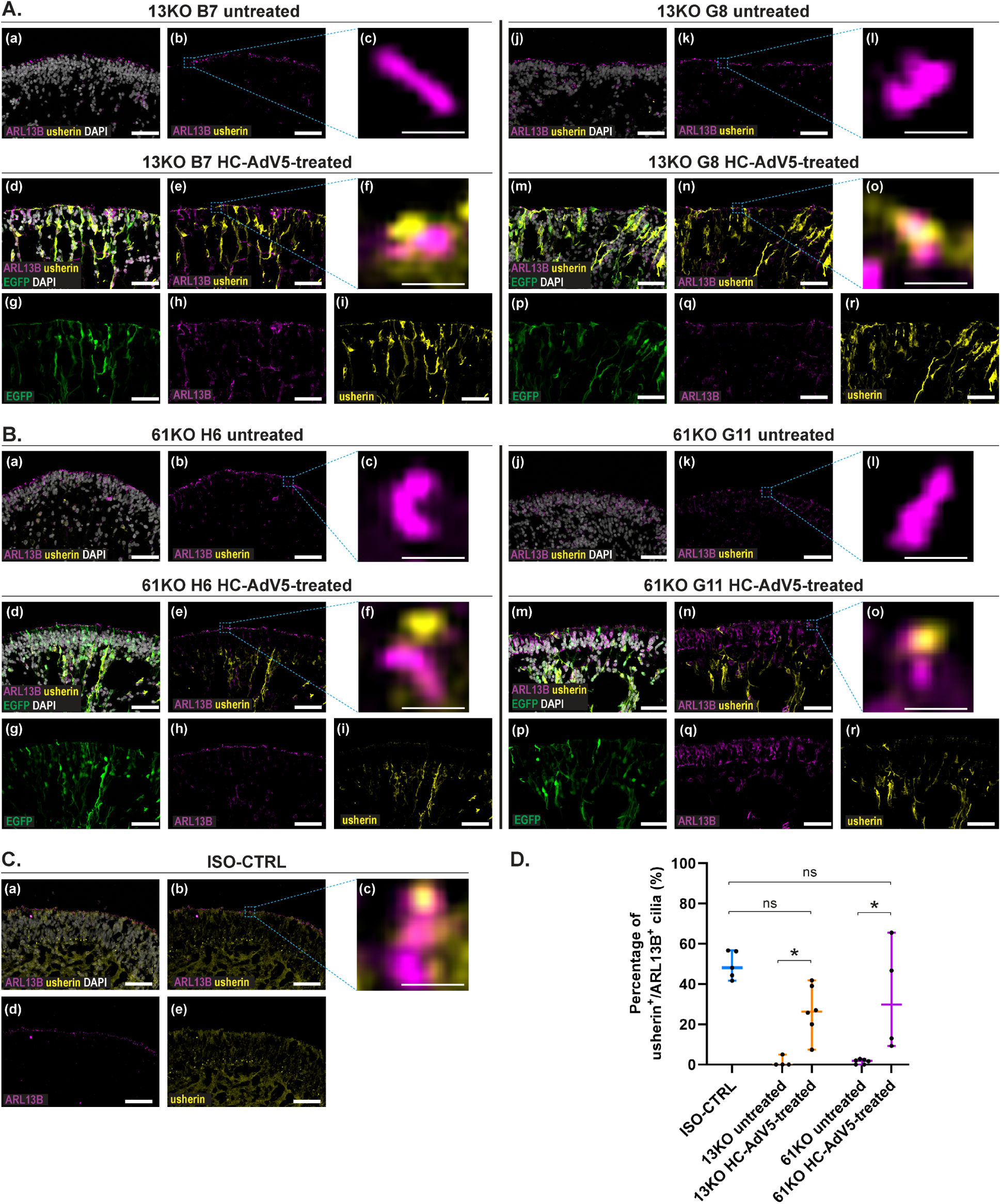
*USH2A* HC-AdV5 transduction of *USH2A* 13KO and 61KO hiPSC-derived retinal organoids shows rescue of usherin at DD240. (A) Representative immunofluorescence Z-stack images of ARL13B (magenta), usherin (yellow), and EGFP (green) in 13KO B7 (left) and 13KO G8 (right) organoids with and without DAPI (grey). Subpanels (a-c) and (j-l) show untreated organoids; subpanels (d-i) and (m-r) show organoids treated with *USH2A* HC-AdV5 and *EGFP* HC-AdV5. Regions outlined by dashed boxes are displayed at higher magnification in the adjacent images. Scale bars in A(a-b, d-e, g-i, j-k, m-n, p-r) are 20 µm; scale bars in A(c, f, l, o) are 1 µm. (B) Representative immunofluorescence Z-stack images of ARL13B (magenta), usherin (yellow), and EGFP (green) in 61KO H6 (left) and 61KO G11 (right) organoids with and without DAPI (grey). Subpanels (a-c) and (j-l) show untreated organoids; subpanels (d-i) and (m-r) show organoids treated with *USH2A* HC-AdV5 and *EGFP* HC-AdV5. Regions outlined by dashed boxes are displayed at higher magnification in the adjacent images. Scale bars in B(a-b, d-e, g-i, j-k, m-n, p-r) are 20 µm; scale bars in B(c, f, l, o) are 1 µm. (C) Representative immunofluorescence Z-stack images of ARL13B (magenta) and usherin (yellow) in ISO-CTRL organoids with and without DAPI (grey). Regions outlined by dashed boxes are displayed at higher magnification in the adjacent image. Scale bars in C(a, b, d, e) are 20 µm; scale bar in C(c) is 1 µm. (A-C) Number of organoids used: ISO-CTRL: *n* = 3, 13KO B7 *n* = 16 (11 untreated, 5 treated), 13KO G8 *n* = 8 (5 untreated, 3 treated), 61KO H6 *n* = 12 (6 untreated, 6 treated), 61KO G11 *n* = 14 (7 untreated, 7 treated), from one differentiation. (D) Quantitative analysis of the percentage of usherin-positive/ARL13B-positive cilia in the photoreceptor layer. Data are presented as median with range. ISO-CTRL: median, 48.1; minimum, 41.7; maximum, 56.7; 13KO untreated: median, 0; minimum, 0; maximum, 5.0; 13KO HC-AdV5-treated: median, 26.4; minimum, 7.4; maximum, 41.9; 61KO untreated: median, 1.8; minimum, 0; maximum, 2.8; 61KO HC-AdV5-treated: median, 29.85; minimum, 9.3; maximum, 65.5. Number of organoids used: ISO-CTRL, n = 5; 13KO untreated, n = 4, 13KO HC-AdV5-treated, n = 6, 61KO untreated, n = 6, 61KO HC-AdV5-treated, n = 4. ISO-CTRL and untreated samples included organoids from at least two independent differentiation batches collected at DD240 and DD225. HC-AdV5-treated samples included organoids from one differentiation batch collected at DD240. 13KO untreated/treated samples represent a pool of B7 and G8 organoids. 61KO untreated/treated samples represent a pool of H6 and G11 organoids. Significance was calculated by Kruskal-Wallis followed by Dunn’s multiple comparisons test and indicated as p < 0.05 (*), ns (not significant). Statistical analysis: 13KO untreated vs 13KO HC-AdV5-treated, *p* = 0.0120; 61KO untreated vs 61KO HC-AdV5-treated, *p* = 0.0167; ISO-CTRL vs 13KO HC-AdV5-treated, *p* = 0.0501; ISO-CTRL vs 61KO HC-AdV5-treated, *p* = 0.4475.

Interestingly, the usherin signal in the *USH2A* KO-treated organoids was observed not only at the connecting cilium, where it physiologically localizes in the ISO-CTRL samples (Figure 4C), but also more broadly throughout the retina, displaying a radial pattern reminiscent of Müller glial cell morphology.

To further assess the extent of usherin restoration at the photoreceptor connecting cilium, we quantified the percentage of ARL13B-positive cilia that also exhibited detectable usherin signal. This analysis revealed a significant increase in the proportion of usherin-positive cilia following *USH2A* HC-AdV5 treatment in both 13KO and 61KO organoids, with the rescued values approaching those observed in the ISO-CTRL samples (Figure 4D).

We next investigated whether the restoration of usherin expression could mediate the relocalization of whirlin and ADGRV1, two interacting components of the photoreceptor periciliary membrane complex. We previously demonstrated that whirlin was below detection levels at the connecting cilium in the *USH2A* 13KO subclones B7 and G8.^11^ Consistent with those findings, here we observed low level of whirlin signal at the photoreceptor connecting cilium of untreated 13KO (B7 and G8) and 61KO (H6 and G11) organoids (Figure 5A(a-c, j-l); Figure 5B(a-c, j-l)).

**Figure 5.**
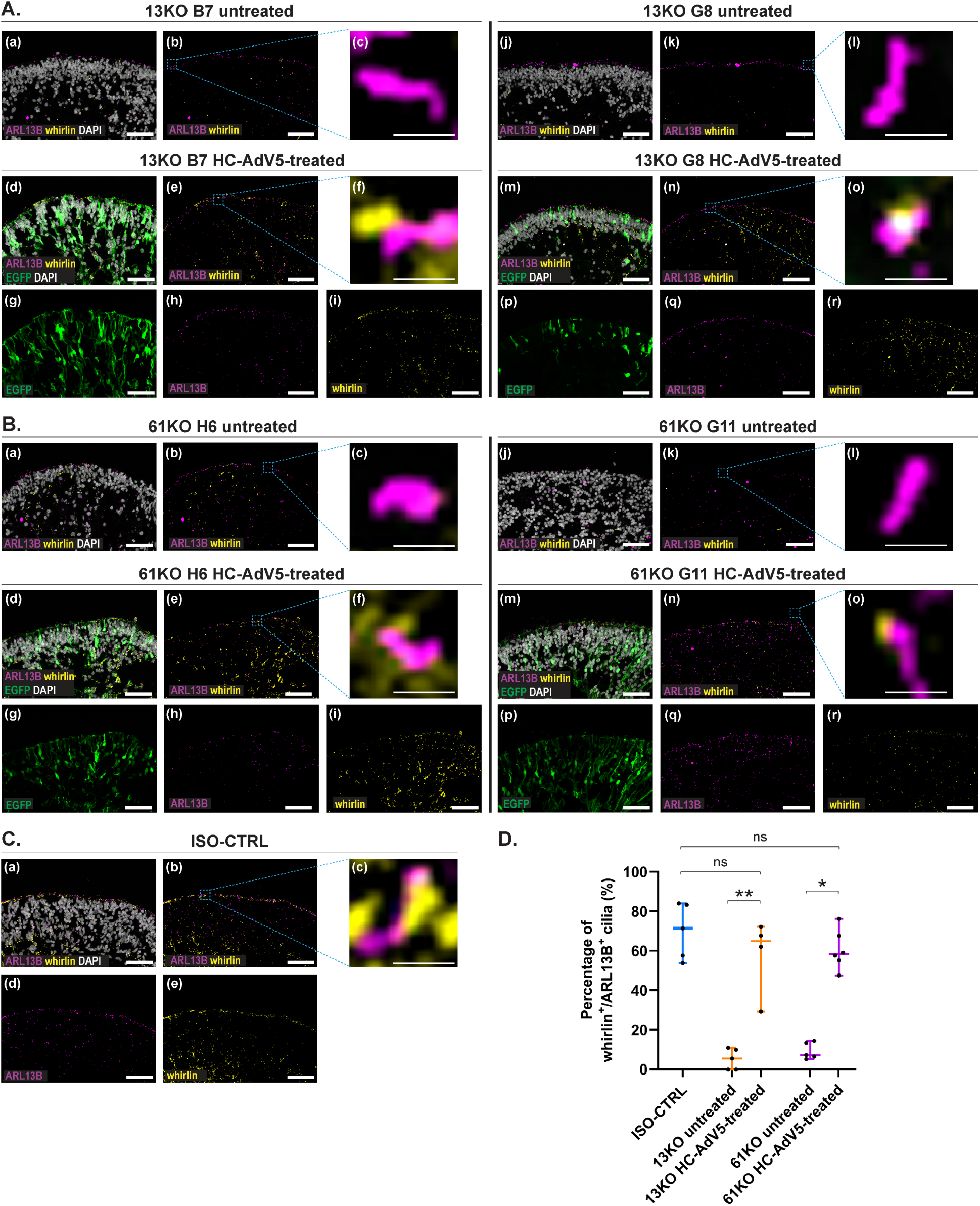
*USH2A* HC-AdV5 transduction of *USH2A* 13KO and 61KO hiPSC-derived retinal organoids shows rescue of whirlin localization at DD240. (A) Representative immunofluorescence Z-stack images of ARL13B (magenta), whirlin (yellow), and EGFP (green) in 13KO B7 (left) and 13KO G8 (right) organoids with and without DAPI (grey). Subpanels (a-c) and (j-l) show untreated organoids; subpanels (d-i) and (m-r) show organoids treated with *USH2A* HC-AdV5 and *EGFP* HC-AdV5. Regions outlined by dashed boxes are displayed at higher magnification in the adjacent images. Scale bars in A(a-b, d-e, g-i, j-k, m-n, p-r) are 20 µm; scale bars in A(c, f, l, o) are 1 µm. (B) Representative immunofluorescence Z-stack images of ARL13B (magenta), whirlin (yellow), and EGFP (green) in 61KO H6 (left) and 61KO G11 (right) organoids with and without DAPI (grey). Subpanels (a-c) and (j-l) show untreated organoids, subpanels (d-i) and (m-r) show organoids treated with *USH2A* HC-AdV5 and *EGFP* HC-AdV5. Regions outlined by dashed boxes are displayed at higher magnification in the adjacent images. Scale bars in B(a-b, d-e, g-i, j-k, m-n, p-r) are 20 µm; scale bars in B(c, f, l, o) are 1 µm. (C) Representative immunofluorescence Z-stack images of ARL13B (magenta) and whirlin (yellow) in ISO-CTRL organoids with and without DAPI (grey). Regions outlined by dashed boxes are displayed at higher magnification in the adjacent image. Scale bars in C(a, b, d, e) are 20 µm; scale bar in C(c) is 1 µm. (A-C) Number of organoids used: ISO-CTRL: *n* = 3, 13KO B7 *n* = 16 (11 untreated, 5 treated), 13KO G8 *n* = 8 (5 untreated, 3 treated), 61KO H6 *n* = 12 (6 untreated, 6 treated), 61KO G11 *n* = 14 (7 untreated, 7 treated), from one differentiation. (D) Quantitative analysis of the percentage of whirlin-positive/ARL13B-positive cilia in the photoreceptor layer. Data are presented as median with range. ISO-CTRL: median, 71.4; minimum, 53.8; maximum, 84; 13KO untreated: median, 5.4; minimum, 0; maximum, 10.8; 13KO HC-AdV5-treated: median, 64.85; minimum, 29.1; maximum, 10.8; 61KO untreated: median, 7.1; minimum, 5; maximum, 14.3; 61KO HC-AdV5-treated: median, 58.35; minimum, 47.5; maximum, 76.2. Number of organoids used: ISO-CTRL, n = 5; 13KO untreated, n = 5, 13KO HC-AdV5-treated, n = 4, 61KO untreated, n = 5, 61KO HC-AdV5-treated, n = 6. ISO-CTRL and untreated samples included organoids from at least two independent differentiation batches collected at DD240 and DD225. HC-AdV5-treated samples included organoids from one differentiation batch collected at DD240. 13KO untreated/treated samples represent a pool of B7 and G8 organoids. 61KO untreated/treated samples represent a pool of H6 and G11 organoids. Significance was calculated by Kruskal-Wallis followed by Dunn’s multiple comparisons test and indicated as p < 0.05 (*), p < 0.01 (**), ns (not significant). Statistical analysis: 13KO untreated vs 13KO HC-AdV5-treated, *p* = 0.0069; 61KO untreated vs 61KO HC-AdV5-treated, *p* = 0.0335; ISO-CTRL vs 13KO HC-AdV5-treated, *p* > 0.9999; ISO-CTRL vs 61KO HC-AdV5-treated, *p* = 0.5511.

To determine whether the decrease in whirlin signal at the connecting cilium reflects an altered subcellular localization rather than reduced protein abundance, we performed Western blot analysis on *USH2A* 61KO retinal organoids at DD225. As previously observed in the 13KO model, whirlin protein was still detectable in *USH2A* 61KO cells (Figure S3), indicating that whirlin protein levels are maintained in both disease models when compared to controls. These observations suggest that disruption of *USH2A* does not result in loss of whirlin expression nor abundance but rather impairs its proper localization within the photoreceptor periciliary membrane complex.

Given that whirlin was retained but mislocalized in the absence of usherin, we next investigated whether transgenic usherin expression could re-establish whirlin localization. Indeed, following *USH2A* HC-AdV5 delivery, both 13KO and 61KO treated organoids showed restoration of whirlin localization at the connecting cilium near ARL13B (Figure 5A(d-I, m-r); Figure 5B(d-l, m-r)), similarly to what was observed in the ISO-CTRL organoids (Figure 5C). Quantitative analysis further confirmed a statistically significant increase in the percentage of whirlin-positive cilia in the photoreceptor layer of both 13KO and 61KO *USH2A* HC-AdV5-treated organoids compared with their respective untreated samples (Figure 5D). The proportion of whirlin-positive signals in the rescued organoids was comparable to that of ISO-CTRL.

A proper localization rescue pattern was also observed for ADGRV1, another key protein of the USH2 complex in photoreceptor cells. Indeed, we confirmed a reduction of ADGRV1 signal at the connecting cilium of untreated *USH2A* 13KO organoids (B7 and G8) and demonstrated its relocation to areas in close proximity to the ARL13B marker upon *USH2A* HC-AdV5 treatment (Figure 6A), suggesting recovery of its proper subcellular distribution. Similarly, in *USH2A* 61KO organoids (H6 and G11), the ADGRV1 signal was only detected near ARL13B in the *USH2A* HC-AdV5-treated cells (Figure 6B), closely resembling the pattern exhibited by ISO-CTRL organoids (Figure 6C). Quantitative analysis confirmed that the percentage of cilia associated with ADGRV1 signals represented a statistically significant increase of this protein in 13KO and 61KO organoids after treatment with *USH2A* HC-AdV5 particles to values closer to the isogenic control ones (Figure 6D).

**Figure 6.**
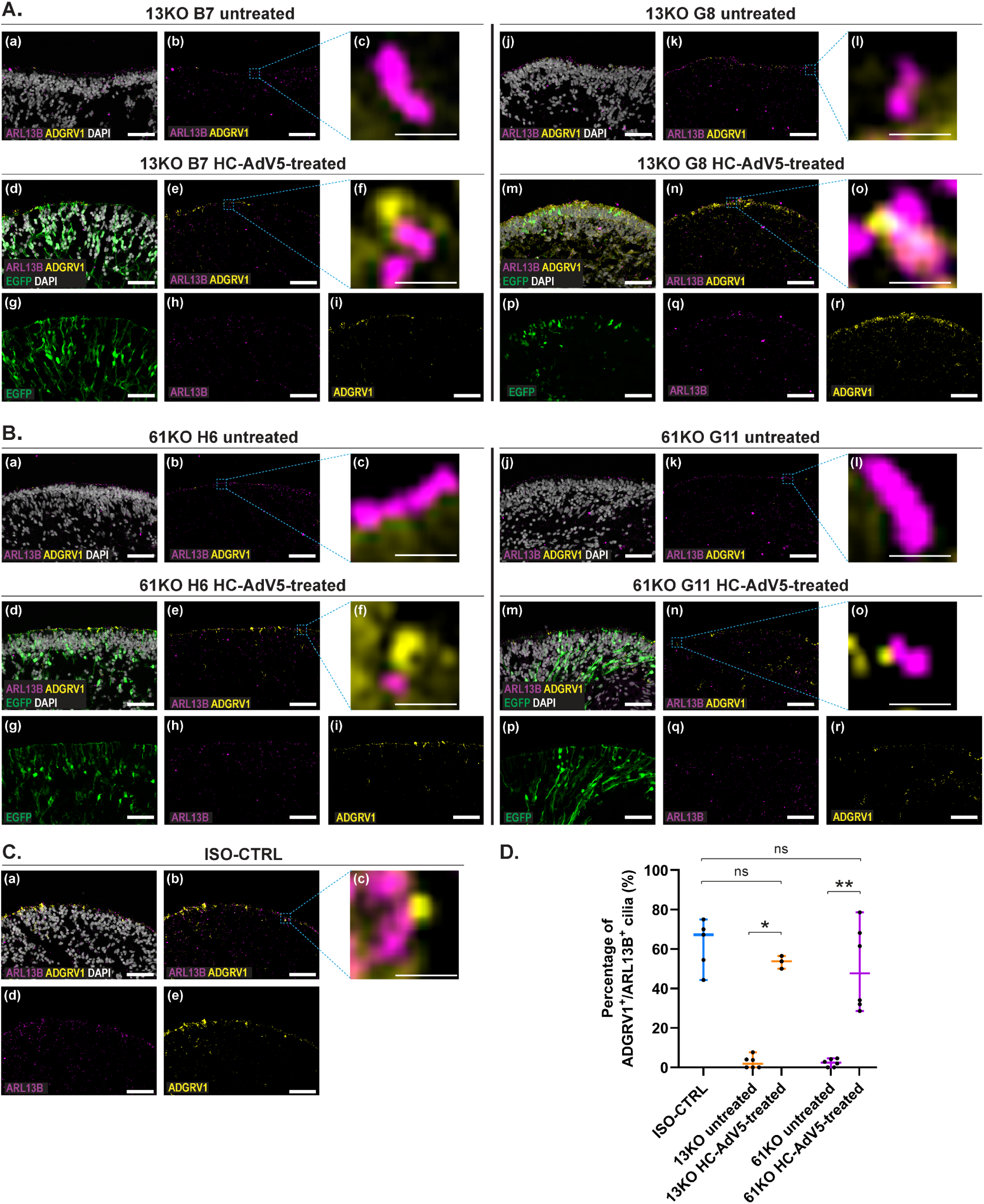
*USH2A* HC-AdV5 transduction of *USH2A* 13KO and 61KO hiPSC-derived retinal organoids shows rescue of ADGRV1 localization at DD240. (A) Representative immunofluorescence Z-stack images of ARL13B (magenta), ADGRV1 (yellow), and EGFP (green) in 13KO B7 (left) and 13KO G8 (right) organoids with and without DAPI (grey). Subpanels (a-c) and (j-l) show untreated organoids; subpanels (d-i) and (m-r) show organoids treated with *USH2A* HC-AdV5 and *EGFP* HC-AdV5. Regions outlined by dashed boxes are displayed at higher magnification in the adjacent images. Scale bars in A(a-b, d-e, g-i, j-k, m-n, p-r) are 20 µm; scale bars in A(c, f, l, o) are 1 µm. (B) Representative immunofluorescence Z-stack images of ARL13B (magenta), ADGRV1 (yellow), and EGFP (green) in 61KO H6 (left) and 61KO G11 (right) organoids with and without DAPI (grey). Subpanels (a-c) and (j-l) show untreated organoids, subpanels (d-i) and (m-r) show organoids treated with *USH2A* HC-AdV5 and *EGFP* HC-AdV5. Regions outlined by dashed boxes are displayed at higher magnification in the adjacent images. Scale bars in B(a-b, d-e, g-i, j-k, m-n, p-r) are 20 µm; scale bars in B(c, f, l, o) are 1 µm. (C) Representative immunofluorescence Z-stack images of ARL13B (magenta) and ADGRV1 (yellow) in ISO-CTRL organoids with and without DAPI (grey). Regions outlined by dashed boxes are displayed at higher magnification in the adjacent image. Scale bars in C(a, b, d, e) are 20 µm; scale bar in C(c) is 1 µm. (A-C) Number of organoids used: ISO-CTRL: *n* = 3, 13KO B7 *n* = 16 (11 untreated, 5 treated), 13KO G8 *n* = 8 (5 untreated, 3 treated), 61KO H6 *n* = 12 (6 untreated, 6 treated), 61KO G11 *n* = 14 (7 untreated, 7 treated), from one differentiation. (D) Quantitative analysis of the percentage of ADGRV1-positive/ARL13B-positive cilia in the photoreceptor layer. Data are presented as median with range. ISO-CTRL: median, 67.3; minimum, 44.4; maximum, 75; 13KO untreated: median, 1.9; minimum, 0; maximum, 7.7; 13KO HC-AdV5-treated: median, 53.8; minimum, 50; maximum, 56.5; 61KO untreated: median, 2.4; minimum, 0; maximum, 4.6; 61KO HC-AdV5-treated: median, 47.8; minimum, 28.6; maximum, 78.6. Number of organoids used: ISO-CTRL, n = 5; 13KO untreated, n = 6, 13KO HC-AdV5-treated, n = 5, 61KO untreated, n = 6, 61KO HC-AdV5-treated, n = 6. ISO-CTRL and untreated samples included organoids from at least two independent differentiation batches collected at DD240 and DD225. HC-AdV5-treated samples included organoids from one differentiation batch collected at DD240. 13KO untreated/treated samples represent a pool of B7 and G8 organoids. 61KO untreated/treated samples represent a pool of H6 and G11 organoids. Significance was calculated by Kruskal-Wallis followed by Dunn’s multiple comparisons test and indicated as p < 0.05 (*), p < 0.01 (**), ns (not significant). Statistical analysis: 13KO untreated vs 13KO HC-AdV5-treated, *p* = 0.0288; 61KO untreated vs 61KO HC-AdV5-treated, *p* = 0.0074; ISO-CTRL vs 13KO HC-AdV5-treated, *p* = 0.6961; ISO-CTRL vs 61KO HC-AdV5-treated, *p* = 0.6345

### Transgene usherin expression is detected in Müller glial cells

To confirm that the detection of usherin was not dependent on the antibody against the usherin C-terminal epitope, we performed co-immunostaining using an anti-HA antibody directed against the HA-tagged HC-AdV5-expressed protein. This analysis showed co-localization of usherin- and HA-specific signals in both *USH2A* HC-AdV5-treated *USH2A* 13KO and 61KO organoids (Figure 7A), revealing consistent detection of recombinant usherin and enabling the use of antibodies from different host species in subsequent experiments.

**Figure 7.**
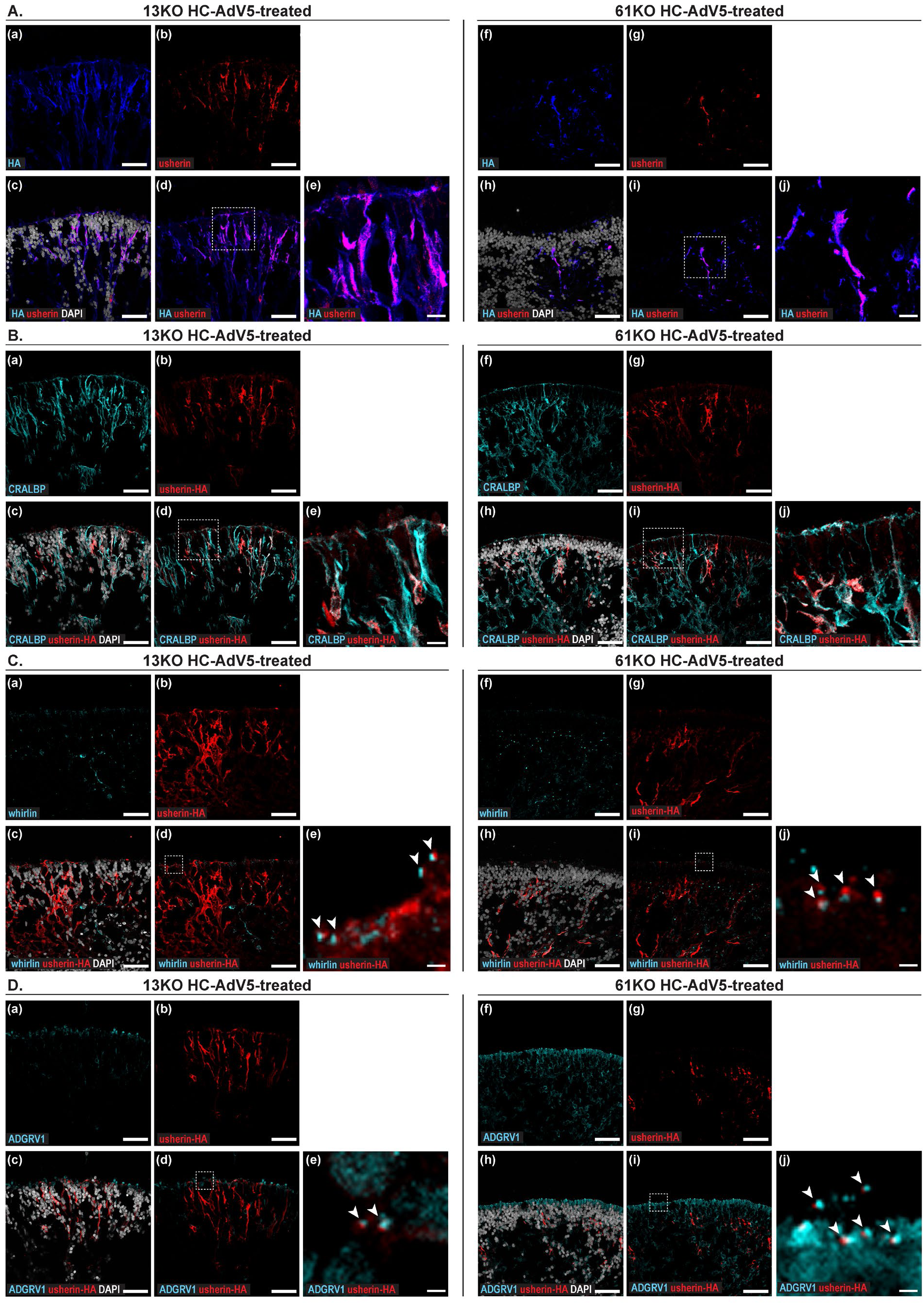
Spatial localization characterization of rescued proteins upon *USH2A* HC-AdV5 transduction of *USH2A* 13KO and 61KO hiPSC-derived retinal organoids. (A) Representative immunofluorescence Z-stack images of HA (blue) and usherin (red) in 13KO (left) and 61KO (right) HC-AdV5-treated organoids with and without DAPI (grey). Regions outlined by dashed boxes are displayed at higher magnification in the adjacent images. Scale bars in A(a-d) and A(f-i) are 20 µm; scale bars in A(e, j) are 10 µm. (B) Representative immunofluorescence Z-stack images of CRALBP (cyan) and HA-tagged usherin (red) in 13KO (left) and 61KO (right) HC-AdV5-treated organoids with and without DAPI (grey). Regions outlined by dashed boxes are displayed at higher magnification in the adjacent images. Scale bars in B(a-d) and B(f-i) are 20 µm; scale bars in B(e, j) are 10 µm. (C) Representative immunofluorescence Z-stack images of whirlin (cyan) and HA-tagged usherin (red) in 13KO (left) and 61KO (right) HC-AdV5-treated organoids with and without DAPI (grey). Regions outlined by dashed boxes are displayed at higher magnification in the adjacent images. Arrows indicate examples of rescued usherin next to whirlin. Scale bars in C(a-d) and C(f-i) are 20 µm; scale bars in C(e, j) are 2 µm. (D) Representative immunofluorescence Z-stack images of ADGRV1 (cyan) and HA-tagged usherin (red) in 13KO (left) and 61KO (right) HC-AdV5-treated organoids with and without DAPI (grey). Regions outlined by dashed boxes are displayed at higher magnification in the adjacent images. Arrows indicate examples of rescued usherin next to ADGRV1. Scale bars in D(a-d) and D(f-i) are 20 µm; scale bars in D(e, j) are 2 µm. (A-D) Number of organoids used: ISO-CTRL: *n* = 3, 13KO B7 *n* = 16 (11 untreated, 5 treated), 13KO G8 *n* = 8 (5 untreated, 3 treated), 61KO H6 *n* = 12 (6 untreated, 6 treated), 61KO G11 *n* = 14 (7 untreated, 7 treated), from one differentiation.

To further investigate whether the usherin-HA signal followed a Müller glial cell-like distribution pattern, we performed immunostaining using anti-cellular retinaldehyde-binding protein (CRALBP) as a Müller glial cell marker together with anti-HA to detect recombinant usherin following *USH2A* HC-AdV5 transduction. Indeed, in addition to the expected localization at the photoreceptor connecting cilium, the HA signal was detected within the Müller glial cells (Figure 7B).

Separately, EGFP expression was detected in both 13KO and 61KO HC-AdV5-treated photoreceptors and Müller glial cells, confirming CMV-driven transgene expression across retinal cell populations (Figure S4), as observed in our previous retinal organoid HC-AdV tropism study.^22^

### USH2A HC-AdV5 restores the close spatial proximity of usherin, whirlin, and ADGRV1 in photoreceptor cells

Finally, we assessed whether the interacting proteins, whirlin and ADGRV1, were restored in proximity to the HA-tagged usherin following *USH2A* HC-AdV5 treatment. Immunofluorescence microscopy analysis detected HA-positive usherin signals adjacent to whirlin (Figure 7C) and ADGRV1 (Figure 7D) signals in the photoreceptor cells of both 13KO and 61KO organoids exposed to *USH2A* HC-AdV5, with occasional co-localization of the respective signals. However, neither whirlin nor ADGRV1 exhibited the radial Müller glia-like distribution pattern observed for the recombinant usherin protein.

## Discussion

The *USH2A* 61KO hiPSC-derived retinal organoids presented in this study and the previously established *USH2A* 13KO model showed that disruption of either exon resulted in reduced amounts of usherin long isoform B and of its interacting partners, ADGRV1 and whirlin, at the photoreceptor connecting cilium. These observations suggest a common pathogenic mechanism involving destabilization of the USH2 usherin-whirlin-ADGRV1 complex, therefore supporting the development of variant-independent therapeutic approaches based on delivery of the full-length *USH2A* coding sequence.

The *USH2A* 61KO organoids did not exhibit alterations in ARL13B ciliary length nor in the outer nuclear thickness, but showed ultrastructural abnormalities in the photoreceptor cells. Differences in sample size, differentiation efficiency, or inter-organoid variability may have influenced the detection of phenotypic changes, warranting further studies. Notably, the comparable distribution of rhodopsin between the photoreceptor outer segments and the outer nuclear layer in mutant and control organoids suggests that *USH2A* disruption does not impair rhodopsin trafficking in photoreceptors with underdeveloped inner and outer segments.

AAV-mediated gene therapy has demonstrated success in inherited retinal dystrophies, with voretigene neparvovec (Luxturna) approved by FDA for *RPE65-*related Leber congenital amaurosis^24^. Although several other strategies are currently under investigation in clinical trials,^25, 26^ the large size of the *USH2A* CDS precludes conventional AAV-mediated approaches. To overcome the limited transgene cargo of AAVs, dual-AAV approaches have been developed to divide large coding sequences into two separate vectors that rely on end-joining recombination, homologous recombination, or RNA trans-splicing to reconstitute a full-length transgene/transcript or, alternatively, depend on protein trans-splicing to reconstitute the full-length protein within co-transduced target cells.^27^ These strategies have shown encouraging preclinical results for large disease genes such as *ABCA4*,^28^ with advances in mRNA trans-splicing^29^ and AAV vectors incorporating translocation linkage strategies^30^ further broadening the repertoire of approaches. Nevertheless, dual-vector systems remain dependent on efficient co-transduction of target cells and precise processing of their derived products, resulting in generally lower efficiency than conventional single-vector AAV delivery.

With a packaging capacity of up to 36-kb, high-capacity adenoviral vectors (HC-AdVs) can deliver large therapeutic genes, eliminating the need for intracellular reassembly of fragmented genetic elements.^31^ Their potential for ocular gene therapy has been investigated in several studies, although most have focused on vector tropism, transduction efficiency, and safety rather than therapeutic gene replacement.^22, 24, 32^ One notable exception is the successful delivery of the 9.4-kb *EYS* CDS to hiPSC-derived retinal organoids from patients with *EYS*-associated retinal dystrophy, where EYS protein localization was restored within 48 hours after transduction at DD200.^33^

Non-viral approaches for the supplementation of large transgenes have also been explored. Scaffold/matrix attachment region (S/MAR)-based episomal DNA vectors mediated the delivery of 15.6-kb human *USH2A* CDS in HEK293 cells, patient-derived fibroblasts, and *ush2a^u507^* zebrafish model, achieving successful protein rescue.^34^

In this study, we provide a proof-of-concept for HC-AdV5-mediated *USH2A* gene supplementation in hiPSC-derived retinal organoids. To our knowledge, this is the first demonstration of viral vector-mediated delivery of a 15.6-kb CDS, expanding beyond previous applications involving large cargos such as the 11.1-kb dystrophin coding sequence.^36^ Here, we successfully delivered the full-length *USH2A* CDS driven from cell-specific regulatory sequences to restore usherin protein expression in both 13KO and 61KO retinal organoids. Importantly, rescue of usherin was accompanied by the re-establishment of ADGRV1 and whirlin at the photoreceptor connecting cilium, suggesting that the amounts of recombinant usherin generated suffice to restore USH2 complex assembly at its proper physiological location. Nevertheless, additional experiments will be required to determine whether recombinant usherin expression and ensuing rescue of whirlin and ADGRV1 localization also recapitulate all functional properties of their respective endogenous complexes.

Previous studies reported that AAV-mediated whirlin replacement in *Whrn*^-/-^ mice restores the mislocalized usherin and ADGRV1 proteins.^37^ Remarkably, our findings reveal that viral vector-mediated usherin rescue can likewise drive the restoration of both ADGRV1 and whirlin at the photoreceptor connecting cilium. Together, these observations support a role for usherin as a central organizer of the USH2 complex localization. Quantitative analysis demonstrated a significant increase in the proportion of cilia positive for usherin, whirlin, and ADGRV1 following *USH2A* HC-AdV5 treatment in both 13KO and 61KO organoids, reaching values closer to those observed in the ISO-CTRL organoids.

Unexpectedly, in this study we detected the recombinant usherin not only in photoreceptor cells, but also within Müller glia cells. Our previous studies have demonstrated that HC-AdVs based on the classical adenoviral type 5 (AdV5) and on a fiber-modified AdV5.F50 version efficiently transduce Müller glial cells in addition to photoreceptors, resulting in transgene delivery to multiple retinal cell types.^22^ Although the *USH2A* CDS was placed under the control of the hGRK1 promoter, widely used to drive photoreceptor-specific expression in AAV vectors,^38–41^ its specificity may not be maintained in HC-AdV-transduced human retinal organoids. We considered whether ectopic hGRK1 promoter activity could arise from regulatory elements within the *USH2A* HC-AdV5 construct. For instance, the SV40 intron positioned upstream of the *USH2A* HC-AdV5 expression cassette may result in high transgene expression in Müller glial cells, potentially by increasing the half-life of *USH2A* mRNA and translation rate.^42^ Alternatively, the vector dose used in this study may have increased transduction of non-target retinal cell populations, thereby revealing low-level hGRK1 promoter activity in cell types beyond photoreceptors. Further studies aimed at optimizing vector dose and promoter specificity, as well as identifying the molecular mechanisms underlying ectopic hGRK1 promoter activity, will be required to improve cell-type-selective targeting.

Importantly, the rescue of usherin and interacting partners at the photoreceptor connecting cilium was here evaluated in single batches of *USH2A* 13KO and 61KO retinal organoids transduced with *USH2A* HC-AdV5. Administration of 4.1 × 10^8^ vp per organoid of *USH2A* HC-AdV5 resulted in robust usherin expression, with the concomitant administration of 0.4 × 10^8^ vp per organoid of *EGFP* HC-AdV5 having no detectable adverse effects in organoid morphology and differentiation capacity, suggesting the absence of overt toxicity associated with *USH2A* HC-AdV5 exposure under the experimental conditions evaluated. Additional rounds of differentiation will be required to confirm the reproducibility and robustness of these findings.

Moreover, future studies should investigate whether HC-AdV5-mediated *USH2A* gene supplementation also rescues the broader transcriptional alterations in retinal homeostasis and activation of innate immune system previously identified in the *USH2A* 13KO Müller glial cells.^11^ Such analyses will provide a more comprehensive assessment of the therapeutic effects of *USH2A* gene supplementation beyond restoration of protein expression and subcellular localization. One batch of untreated and *USH2A* HC-AdV5-treated 13KO organoids was processed at DD240 for single-cell RNA sequencing. However, the resulting single-cell suspensions did not meet the required quality criteria for downstream analysis due to low cell viability. Therefore, this experiment will need to be performed in future studies.

Despite the removal of all viral coding sequences, HC-AdVs retain adenoviral capsid proteins that can trigger immune responses.^43^ Consistent with this concern, subretinal delivery of HC-AdV5 vectors expressing *EGFP* in rat retinas has been reported to induce an acute inflammatory response, characterized by vitreoretinal infiltration of Iba1-positive microglial cells, increased expression of inflammatory markers, and subsequent photoreceptor loss.^24^ In contrast, in hiPSC-derived retinal organoids, delivery of *EGFP* HC-AdV5 was not associated with evidence of reactive gliosis or photoreceptor cell death, although an increase in the outer nuclear layer thickness was measured at 110 days post-transduction.^22^ Similarly, subretinal administration of HC-AdV vectors expressing the *LacZ* reporter in immunocompetent rats resulted in stable transgene expression without significant morphological changes or visible signs of inflammation.^20^ These findings suggest that the safety profile of HC-AdV delivery may depend on the experimental model, vector characteristics, and conditions of administration, highlighting the need for further assessment *in vivo*.^20^

HC-AdV-based retinal gene therapies will profit from deepening our knowledge about vector particle-cell interactions that induce pro-inflammatory cytokine release or activate intracellular innate immune sensors.^44^ The resulting findings should guide strategies to, for instance, transiently modulate these signaling pathways to enhance therapeutic efficacy and safety. Moreover, further progress in capsid engineering to achieve strict cell type-specific transduction and, at the same time, mitigate immune cell delivery (e.g., tissue-resident antigen-presenting cells), is required.^45^

In this study, we demonstrated that HC-AdV5-mediated *USH2A* gene supplementation in *USH2A*-mutant retinal organoids restores usherin expression and re-establishes the physiological localization of whirlin and ADGRV1 at the photoreceptor connecting cilium. These findings provide proof-of-concept for viral-mediated full-length *USH2A* gene supplementation as a potential therapeutic strategy for *USH2A*-associated retinal disease, warranting further studies to assess its safety, efficacy, and feasibility for patient treatment.

## Materials and Methods

### hiPSC genome editing by CRISPR/Cas9

The Integrated DNA Technologies (IDT) CRISPR design tool (https://eu.idtdna.com/; access date, June 2023) was used to select a gRNA targeting exon-61 of *USH2A.* A single-stranded oligodeoxynucleotide (ssODN) HDR template (IDT, Madison, WI, USA) was designed to introduce a CATG sequence, generating a downstream premature stop codon and a BspHI restriction site, to facilitate clone screening. Genome editing was performed using the Neon Transfection System (ThermoFisher Scientific, Waltham, MA, USA) as previously described.^11^ Edited 61KO clones were validated by Sanger sequencing. Top five off-target sites were predicted using the Integrated DNA Technologies (IDT) CRISPR-Cas9 guide RNA design checker tool (https://eu.idtdna.com/; access date, June 2023). Additional information on hiPSC lines and reagents is provided in Tables S1-S3.

### Cell Culture and Retinal Organoid Differentiation

hiPSCs were cultured under standard conditions and differentiated into retinal organoids according to a previously established protocol.^11, 22^ Briefly, embryoid bodies were generated in agarose micro-molds and sequentially transitioned through neural induction and retinal maturation media supplemented with SAG (Selleck Chemicals, Houston, TX, USA), retinoic acid (Merck, Schiphol-Rijk, The Netherlands), and DAPT (Selleck Chemicals, Houston, TX, USA). Retinal organoids were maintained in long-term culture until collection. Additional reagent information can be found in Table S3.

### Immunofluorescence analysis

Retinal organoids were fixed with 4% paraformaldehyde in PBS for 20 minutes. After a quick wash in PBS, they were incubated first in 15% sucrose PBS solution for 30 minutes at RT, and later in a 30% sucrose PBS solution for at least 1 h at RT. The organoids were embedded in Tissue-Tek O.C.T. Compound (Sakura Finetek Europe, Alphen aan den Rijn, The Netherlands) and cryosectioned at 8 μm thickness using a Leica CM1900 cryostat (Leica Microsystems, Wetzlar, Germany). Immunofluorescence analysis was performed as previously described.^11^ Images were acquired using a Leica TCS SP8 confocal microscope (Leica Microsystems, Wetzlar, Germany) with Leica Application suite X software (v3.7.0.20979). Antibodies used in this study are listed in Table S4. EGFP signal was detected by direct fluorescence microscopy using a 488 nm excitation laser.

### Transmission electron microscopy analysis

Retinal organoids were fixed by adding an equal volume of 3% glutaraldehyde in 0.2 M cacodylate buffer to the culture medium, resulting in a final concentration of 1.5% glutaraldehyde in 0.1 M cacodylate buffer, and incubated for 1 h at room temperature. Samples were rinsed three times with 0.1 M cacodylate buffer and post-fixed in 1% osmium tetroxide (OsO₄) and 1.5% potassium ferricyanide in 0.1 M cacodylate buffer for 1 h on ice. Following three additional washes in 0.1 M cacodylate buffer, samples were dehydrated through graded ethanol and acetone series and infiltrated with increasing concentrations of EPON resin (LX112; Ladd Research Industries, Essex Junction, VT, USA), followed by embedding in 100% EPON. Embedded organoids were polymerized at 70°C for 48 h. Ultrathin sections (90 nm) were prepared using a Reichert Ultracut S ultramicrotome (Leica Microsystems, Wetzlar, Germany), stained with uranyl acetate and lead citrate, and imaged using a Tecnai T12 Twin transmission electron microscope (Thermo Fisher Scientific, Eindhoven, The Netherlands) operating at 120 kV and equipped with a OneView camera (Gatan). Overlapping images were acquired and stitched into composite images as previously described.^46^

### Quantification and Statistical analysis

Three representative images per organoid were acquired at 40× magnification and used for quantification of three independent rounds of differentiation on Fiji ImageJ (v2.17). For usherin, whirlin, and ADGRV1 protein rescue, quantification was instead performed on a single batch of HC-AdV5-treated retinal organoids and at least two differentiation batches (DD225 and DD240) of ISO-CTRL, 13KO, and 61KO untreated organoids. Data were plotted as medians and ranges on GraphPad Prism (v10.6.1). The number of tested organoids and the minimum and maximum values of ranges are indicated in the figure legends. Statistical significance was calculated using the Kruskal–Wallis test followed by Dunn’s multiple comparisons test, and was indicated as p < 0.05 (*), p < 0.01 (**), and ns (not significant).

### Production and Characterization of High-Capacity Adenoviral Vectors

A 33.3-kb plasmid (pGLAd) carrying a 30.3-kb HC-AdV molecular clone insert containing the hGRK1 promoter and a codon-optimized CDS for the human full-length usherin protein, was customer-designed at Leiden University Medical Center, without the use of VectorBuilder software. The customer-built *USH2A* plasmid was then used to generate HC-AdV5 vector particles. Ultra-purified *USH2A* HC-AdV5 particles were produced by VectorBuilder (Chicago, IL, USA). *EGFP* HC-AdV5 particles, were obtained from the pGLAd[Exp]-CMV>*EGFP* plasmid (VectorBuilder, Chicago, IL, USA) and used as control. Adenoviral vector particles were stored in GTS buffer (2.5% glycerol, 20 mM Tris, 25 mM NaCl, pH 8.0) at −80 °C until use. Vector titers, determined by optical absorbance at 260 nm, were 1.4 x 10^11^ viral particles (vp)/mL for *USH2A* HC-AdV5 and 2.7 x 10^11^ vp/ml for *EGFP* HC-AdV5. Vector preparations were tested by the manufacturer and certified free of microbial and mycoplasma contamination.

### High-capacity Adenoviral Vector Transduction of Human iPSC-Derived Retinal Organoids

At DD134, retinal organoids were transferred to agarose-coated 96-well plates. At DD135, individual organoids were transduced with a mix of HC-AdV5 particles diluted in RLM2 medium and incubated for 8 h at 37 °C and 5% CO₂. Pipette tips were pre-rinsed with 0.001% Poloxamer 188 in DPBS prior to pipetting vector solutions to reduce non-specific adsorption of viral particles to plastic surfaces. Each organoid received 4.1 × 10^8^ vp of *USH2A* HC-AdV5 and 0.4 × 10^8^ vp of *EGFP* HC-AdV5 to monitor transduction efficiency. Following the 8-hour incubation period, the organoids were transferred to a freshly prepared 48-well agarose plate and maintained in RLM2 medium under standard culture conditions until collection at DD240. A part of the organoids from each line were left untreated to serve as controls for phenotype comparisons.

## Data availability

The supporting data of this study can be obtained from the corresponding author upon reasonable request.

## Acknowledgments

Uitzicht 2021-11 to J.W.; Uitzicht 2023-05 to J.W.; Uitzicht 2024-3 to E.v.D. and J.W.; Uitzicht 2025-11 to E.v.D. and J.W.: Stichting UsherSyndroom, Stichting Oogfonds, Rotterdamse Stichting Blindenbelangen, Landelijke Stichting voor Blinden en Slechtzienden, Stichting Retina Nederland Fonds, and Stichting Blindenhulp.

The authors thank Dr. Jun Yang for providing the antibodies against usherin isoform B and ADGRV1, and their team members for valuable discussions and advice.

## Author contributions

Conceptualization, R.V., M.A.F.V.G., and J.W.; design phGRK1:SV40-intron>*USH2A*-3xHA:WPRE3.HSV-TK-pA plasmid, J.W.; methodology, R.V., X.L., A.A.M., and R.I.K.; validation, R.V.; formal analysis, R.V.; investigation, R.V., A.A.M., and R.I.K.; writing—original draft preparation, R.V.; writing—review and editing, R.V. and J.W.; visualization, R.V.; supervision, J.W.; project administration, J.W.; funding acquisition, E.v.D. and J.W. All authors have read and agreed to the published version of the manuscript.

