## Supplemental files for "*USH2A* High-Capacity Adenoviral Vector Restores usherin, whirlin, and ADGRV1 in Photoreceptor Cells of Retinal Organoids"

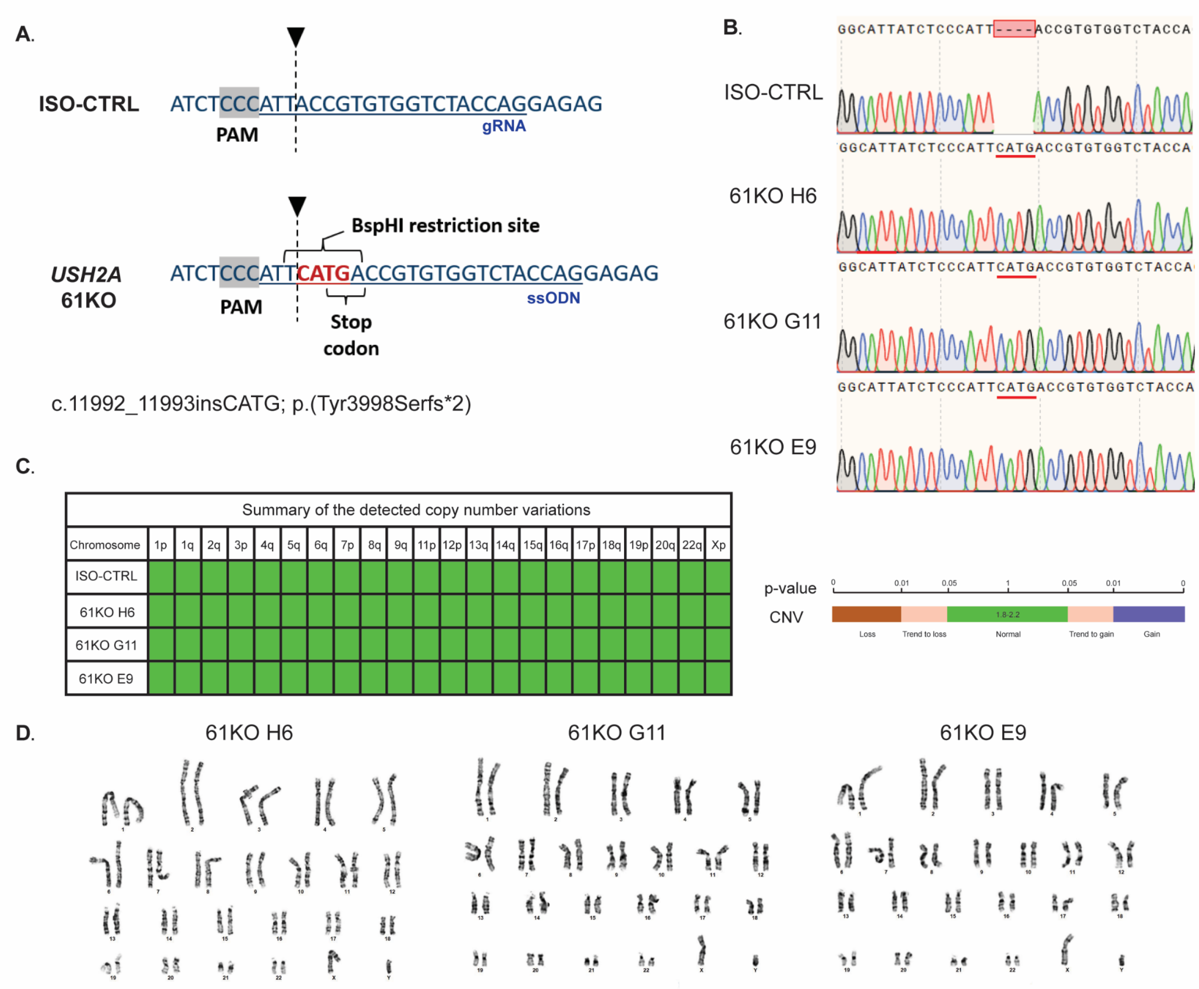


E.

| Short tandem repeat profile | | | | | | | | | | | |
|---|---|---|---|---|---|---|---|---|---|---|---|
| Markers | CSF1PO | D1S1656 | D2S1338 | D2S441 | D3S1358 | D5S818 | D7S820 | D8S1179 | D10S1248 | D12S391 | D13S317 |
| ISO-CTRL | 11 | 16.3,17.3 | 19,20 | 12,14 | 17,18 | 12,13 | 9,10 | 14 | 13,15 | 19,22 | 11,13 |
| 61KO H6 | 11 | 16.3,17.3 | 19,20 | 12,14 | 17,18 | 12,13 | 9,10 | 14 | 13,15 | 19,22 | 11,13 |
| 61KO G11 | 11 | 16.3,17.3 | 19,20 | 12,14 | 17,18 | 12,13 | 9,10 | 14 | 13,15 | 19,22 | 11,13 |
| 61KO E9 | 11 | 16.3,17.3 | 19,20 | 12,14 | 17,18 | 12,13 | 9,10 | 14 | 13,15 | 19,22 | 11,13 |

| Markers | D16S539 | D18S51 | D19S433 | D21S11 | D22S1045 | FGA | Penta D | Penta E | TH01 | TPOX | vWA |
|---|---|---|---|---|---|---|---|---|---|---|---|
| ISO-CTRL | 8,15 | 13 | 12 | 27,30 | 16 | 24 | 10,12 | 5,12 | 7,10 | 8 | 16,18 |
| 61KO H6 | 8,15 | 13 | 12 | 27,30 | 16 | 24 | 10,12 | 5,12 | 7,10 | 8 | 16,18 |
| 61KO G11 | 8,15 | 13 | 12 | 27,30 | 16 | 24 | 10,12 | 5,12 | 7,10 | 8 | 16,18 |
| 61KO E9 | 8,15 | 13 | 12 | 27,30 | 16 | 24 | 10,12 | 5,12 | 7,10 | 8 | 16,18 |

**Figure S1: Generation and characterization of *USH2A* 61KO hiPSCs.** (A) Design of the gene editing strategy in *USH2A* exon-61. The single stranded oligodeoxynucleotide (ssODN) mediates the insertion of the CATG sequence upstream the PAM of the selected gRNA, resulting in an in-frame premature stop codon and the formation of a BspHI restriction site to facilitate screening of the edited clones. (B) Sanger sequencing validation of the genetic editing of clones 61KO H6, G11, and E9. (A, C, D, E) Restriction enzyme BspHI (BspHI); guide RNA (gRNA); protospacer adjacent motif (PAM) (A), copy number variation (CNV) (C), karyotyping (D), and short tandem repeat (STR) (E) analyses of *USH2A* 61KO clones.

**Supplemental**

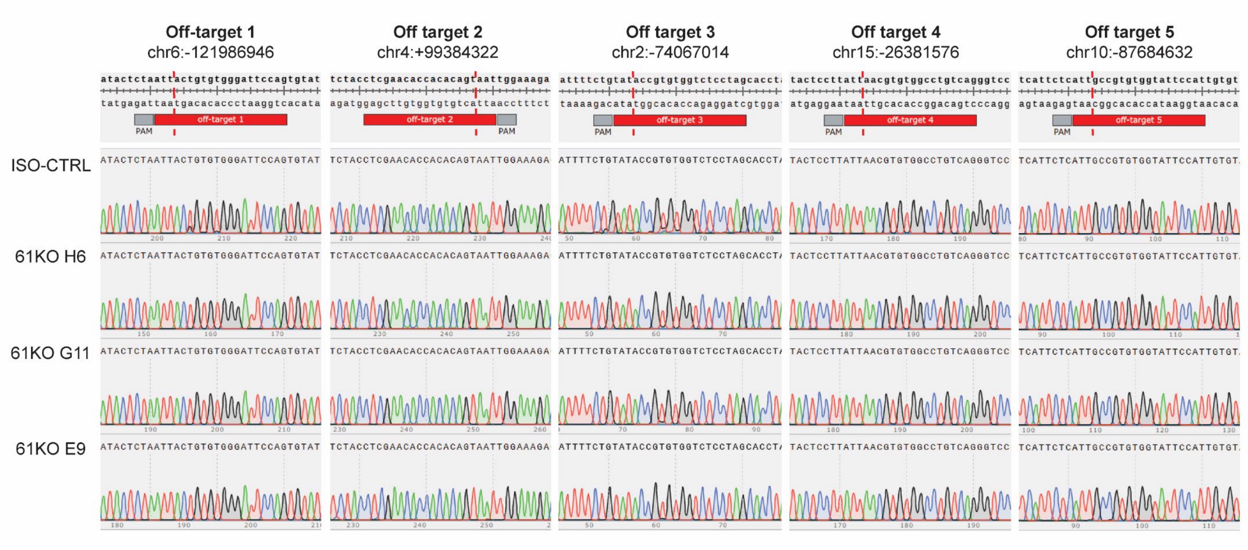


**Figure S2: Off-target mutagenesis analysis of *USH2A* 61KO hiPSCs.** Analysis of the top 5 off-target candidates associated with the gRNA mediating the CRISPR-Cas9 gene editing in *USH2A* exon-61 as predicted using the CRISPR-Cas9 guide RNA design checker tool provided by Integrated DNA Technologies (IDT) . Chr6:-121986946: chromosome 6, position -121986946; chr4:+99384322: chromosome 4, position +99384322; chr2:-74067014: chromosome 2, position -74067014; chr15:-26381576: chromosome 15, position -26381576; chr10:-87684632: chromosome 10, position -87684632.

**Supplemental**

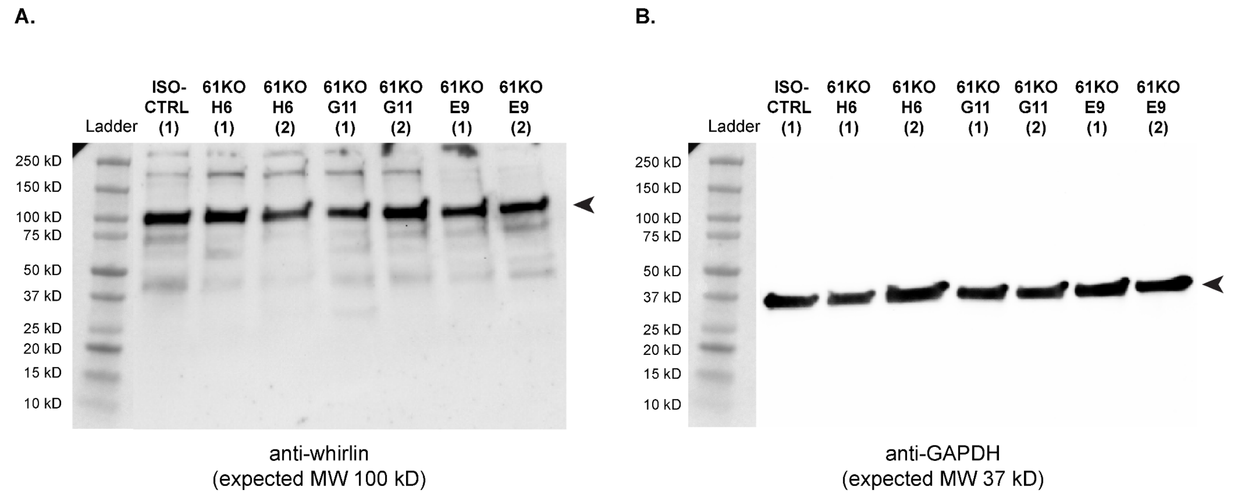


**Figure S3: Western blot analysis of whirlin in control and *USH2A* 61KO retinal organoids at DD225.** (A) Detection of whirlin in lysates from ISO-CTRL (n = 1), *USH2A* 61KO H6 (n = 2), *USH2A* 61KO G11 (n = 2), and *USH2A* 61KO E9 (n = 2) retinal organoids. (B) Detection of glyceraldehyde 3-phosphate dehydrogenase (GAPDH) in lysates from ISO-CTRL (n = 1), *USH2A* 61KO H6 (n = 2), *USH2A* 61KO G11 (n = 2), and *USH2A* 61KO E9 (n = 2) retinal organoids. (A, B) Each lane was loaded with 20 µg of protein lysate in a final volume of 40 µl. Arrowheads point at the expected band height for whirlin (A) and GAPDH (B).

### Supplemental

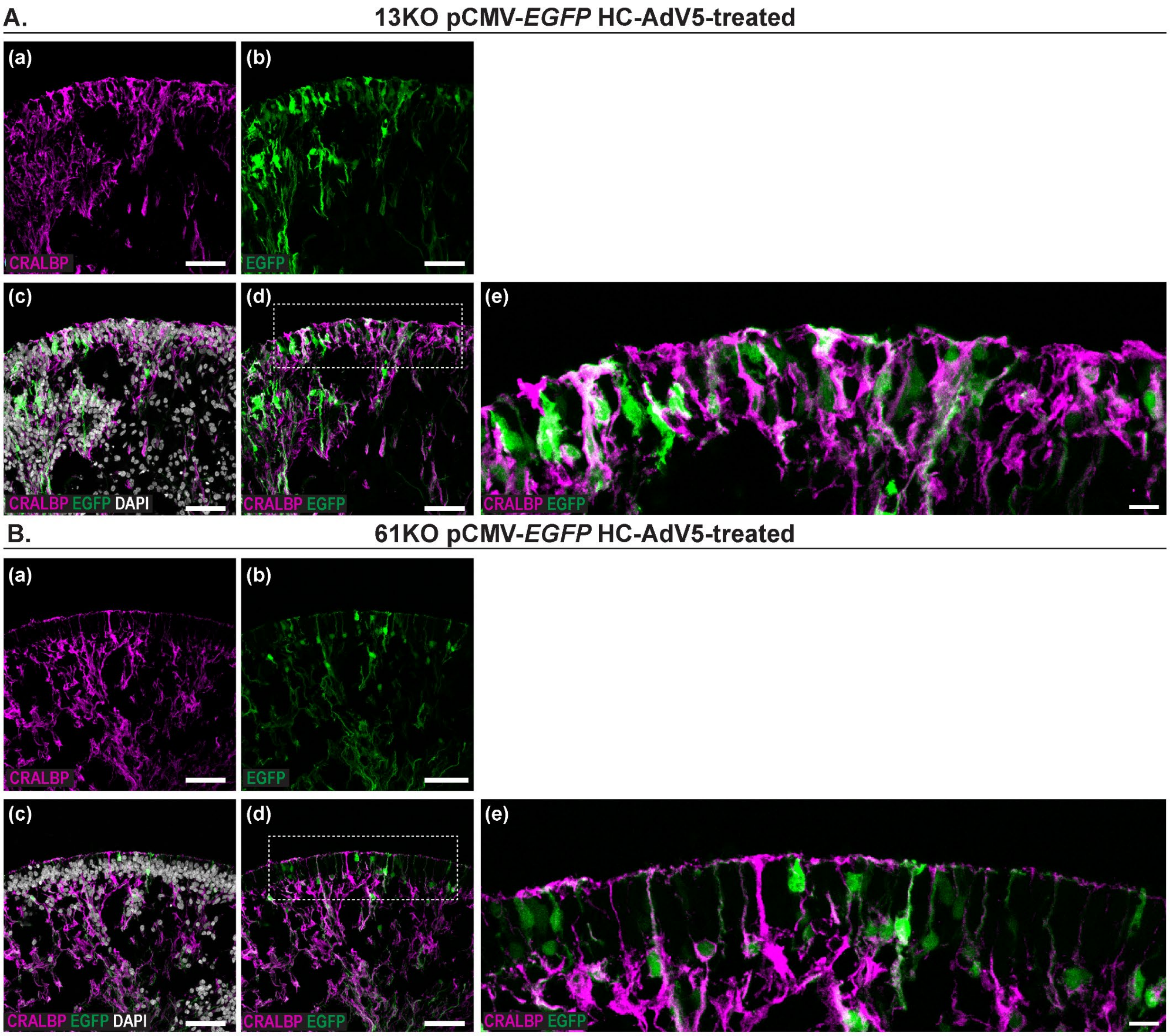


**Figure S4: *EGFP* HC-AdV5 transduction of *USH2A* 13KO and 61KO hiPSC-derived retinal organoids reveals expression of EGFP in both photoreceptors and MGCs at DD240.** (A-B) Representative immunofluorescence Z-stack images of CRALBP (magenta) and EGFP (green) in 13KO (A) and 61KO (B) pCMV-*EGFP* HC-AdV5-treated organoids with and without DAPI (grey). pCMV, CMV promoter. Regions outlined by dashed boxes are displayed at higher magnification in the adjacent image. Scale bars in A(a-d) and B(a-d) are 20 µm; scale bars in A(e) and B(e) are 10 µm.

**Supplemental**

**Table S1: hiPSC line information.**

| Line name | Description | Gender |
|---|---|---|
| **LUMC0004iCTRL10 = ISO-CTRL (hPSCreg name: LUMCi029-B)** | Control parental hiPSC line | Male |
| **LUMC0004iCTRL10_*USH2A*$^{13KO}$ CLB7 = 13KO B7** | It has a stop codon in exon-13 of *USH2A*. c.2553_2554insTAGT, p.(Thr852*) | Male |
| **LUMC0004iCTRL10_*USH2A*$^{13KO}$ CLG8 = 13KO G8** | It has a stop codon in exon-13 of *USH2A*. c.2553_2554insTAGT, p.(Thr852*) | Male |
| **LUMC0004iCTRL10_*USH2A61*$^{KO}$ CLH6 = 61KO H6** | It has a stop codon in exon-61 of *USH2A*. c.11992_11993insCATG, p.(Tyr3998Serfs*2) | Male |
| **LUMC0004iCTRL10_*USH2A61*$^{KO}$ CLG11 = 61KO G11** | It has a stop codon in exon-61 of *USH2A*. c.11992_11993insCATG, p.(Tyr3998Serfs*2) | Male |
| **LUMC0004iCTRL10_*USH2A61*$^{KO}$ CLE9 = 61KO E9** | It has a stop codon in exon-61 of *USH2A*. c.11992_11993insCATG, p.(Tyr3998Serfs*2) | Male |

Information on the hiPSC lines used in this study.

**Supplemental**

**Table S2: Sequences of the crRNA and donor template (ssODN) used to generate the *USH2A* 61KO hiPSC lines and chosen primers to validate the genomic mutation and absence of off-target mutagenesis.**

| **Name** | **Sequence (5'-3')** |
|---|---|
| *USH2A* exon-61 crRNA | ATTACCGTGTGGTCTACCAG |
| *USH2A* ssODN exon-61 | /AlT-R-HDR1/G*A*TTTTCCAGCTCCTTGGGCTCAAGCCACGAGTGCTCATTCAGTTCTGTTGAATTGGACAAAGCCAGAATCTCCCAATGGCATTATCTCCCATTCATGACCGTGTGGTCTACCAGGAGAGACCCGACGATCCTACATTTAACAGCCCTACCGTGCATGCTTTCACAGTGAAGGTAAGACCCTTTAGAAAA*G*T/AlT-R-HDR2/ |
| *USH2A* exon-61 Fw primer | TGAAGTGTGCAGCTGTCACT |
| *USH2A* exon-61 Rv primer | AGCCCTAAGTGAAGAAAAATGAGA |
| Off-target 1 (chr6:-121986946) Fw primer | TCCAGCACCTCAGAAAGCTC |
| Off-target 1 (chr6:-121986946) Rv primer | TCCGGAGTAGCTGGGACTAC |
| Off-target 2 (chr4:+99384322) Fw primer | CAGGGTACCACGTGCAAAGT |

| | |
|---|---|
| Off-target 2 (chr4:+99384322) Rv primer | ACAGGCAACTCTGACCCCAT |
| Off-target 3 (chr2:-74067014) Fw primer | TTCCCAGCATAAGACCCTGC |
| Off-target 3 (chr2:-74067014) Rv primer | CGGAAACCTGGGCAACTACA |
| Off-target 4 (chr15:-26381576) Fw primer | GTAAGCTCAGGCCACTCAGG |
| Off-target 4 (chr15:-26381576) Rv primer | GAGGCCAGAGACGAACTTCC |
| Off-target 5 (chr10:-87684632) Fw primer | TCCAACACAGGATGTCTGCC |
| Off-target 5 (chr10:-87684632) Rv primer | TCAAGGTCTGGCTCAACCAC |

Forward (Fw) and reverse (Rv) primers used to confirm the CRISPR/Cas9-mediated insertion of CATG in *USH2A* exon-61 and the absence of the top 5 off-targets as predicted by the online tools provided by Integrated DNA Technologies (IDT).

**Supplemental**

**Table S3: List of materials used in this study.**

| **Materials** | **Source** | **Identifier** |
|---|---|---|
| **tracrRNA** | IDT | 1072532 |
| **Nuclease-Free Duplex Buffer** | IDT | 11-01-03-01 |
| **SpCas9 Nuclease V3** | IDT | 1081058 |
| **Neon™ Transfection System kit** | Invitrogen | MPK10096 |
| **Neon™ Transfection System** | Invitrogen | MPK1025 |
| **Matrigel hESC-Qualified Matrix** | Corning | 354277 |
| **mTeSR plus medium** | STEMCELL Technologies | 100-0276 |
| **Accumax** | STEMCELL Technologies | 07921 |
| **Gentle Cell Dissociation reagent** | STEMCELL Technologies | 100-0485 |
| **CloneR** | STEMCELL Technologies | 05888 |
| **Cryostor** | STEMCELL Technologies | 07930 |
| **DMEM/F12** | Life Technologies | 11320074 |
| **DMEM(1X) + GlutaMAX** | ThermoScientific | 10569010 |
| **Fasudil HCl** | Focus Biomolecules | 10-2137 |
| **40 µm cell strainer** | PluriSelect | 43-10040-40 |

**Supplemental**

| | | |
|---|---|---|
| **BspHI** | ThermoScientific | ER1281 |
| **Blebbistatin** | Abcam | ab120425 |
| **Micro-molds** | Merck | Z764000-6EA |
| **MEM NEAA 100X** | Life technologies | 11140-035 |
| **Taurine** | Merck | T0625 |
| **Neurocult SM1 50X** | STEMCELL Technologies | 05711 |
| **N2 supplement 100X** | Life technologies | 17502048 |
| **Heparin** | Merck | H-9399 |
| **Smoothened agonist (SAG)** | Selleck Chemicals | S7779 |
| **Gamma secretase inhibitor IX (DAPT)** | Selleck Chemicals | S2215 |
| **Fetal Bovine Serum** | Serana | S-FBS-CO-015 |
| **Poloxamer 188** | Merck | P5556 |
| **Retinoic acid** | Merck | R-2625 |
| **Antibiotic-antimycotic 100X** | Merck | A5955 |
| **Tissue-Tek O.C.T. Compound** | Sakura Finetek | 4583 |

**Supplemental**

| | | |
|---|---|---|
| **Vectashield Antifade Mounting Medium** | Vector Laboratories | H-1800-10 |
| **DPBS** | ThermoScientific | 14040117 |
| **pGLAd[Exp]-CMV>*EGFP, EGFP* HC-AdV5** | VectorBuilder | #GLAdCP(VB010000-9400ggg) |

**Supplemental**

**Table S4: List of antibodies used in this study.**

| Antibody | Dilution | Source | Identifier |
|---|---|---|---|
| **Anti-Rhodopsin** | 1:500 | Millipore | MAB5356 |
| **Anti-F-actin** | 1:300 | Thermo Fisher Scientific | r-415 |
| **Anti-usherin (C-terminal)** | 1:2500 | Gift by Dr. Yang | NA |
| **Anti-ADGRV1** | 1:7500 | Gift by Dr. Yang | NA |
| **Anti-whirlin** | 1:400 (IF) 1:1000 (WB) | Proteintech | 25881-1-AP |
| **Anti-ARL13B** | 1:200 | Proteintech | 66739-1-Ig |
| **Anti-HA (mouse)** | 1:500 | Biolegend | 901513 |
| **Anti-HA (rabbit)** | 1:500 | Proteintech | 51064-2-AP |
| **Anti-CRALBP** | 1:200 | Invitrogen | MA1-813 |
| **Anti-GAPDH** | 1:1000 | Proteintech | 60004-1-Ig |
| **Goat anti-rabbit IgG (H+L) Highly Cross-Adsorbed Secondary Antibody, Alexa Fluor 488** | 1:1000 | Invitrogen | A-11034 |
| **Goat anti-mouse IgG H&L, Alexa Fluor 555** | 1:1000 | Abcam | ab150118 |

| | | | |
|---|---|---|---|
| **Cy™3 AffiniPure® Goat Anti-Rabbit IgG (H+L)** | 1:1000 | Jackson ImmunoResearch | 111-165-045 |
| **Goat anti-mouse IgG H&L, Alexa Fluor 647** | 1:1000 | Abcam | ab150119 |
| **Anti-rabbit-IgG-HRP** | 1:5000 | Santa Cruz Biotechnology | sc-2357 |
| **Anti-mouse-IgGκ BP-HRP** | 1:5000 | Santa Cruz Biotechnology | sc-516102 |

Product information about the antibodies used in immunofluorescence and Western Blot analyses.

**Supplemental**

**Supplemental methods – Western blot analysis**

Protein lysates were obtained from ISO-CTRL and *USH2A* 61KO organoids as previously described.[10] Following wet overnight transfer, the polyvinylidene fluoride (PVDF) membrane was blocked with 5% milk in PBST and then incubated overnight at 4 °C with anti-whirlin (1:1000) diluted in 5% milk-PBST. The blot was imaged with the Chemidoc system (Bio-Rad, Hercules, CA, USA). After stripping, GAPDH (glyceraldehyde 3-phosphate dehydrogenase) immunodetection was performed to control for loading variability.
